# *specificity*: an R package for analysis of feature specificity to environmental and higher dimensional variables, applied to microbiome species data

**DOI:** 10.1101/2021.11.06.467582

**Authors:** John L. Darcy, Anthony S. Amend, Sean O. I. Swift, Pacifica S. Sommers, Catherine A. Lozupone

## Abstract

**Background:** Understanding the factors that influence microbes’ environmental distributions is important for determining drivers of microbial community composition. These include environmental variables like temperature and pH, and higher-dimensional variables like geographic distance and host species phylogeny. In microbial ecology, “specificity” is often described in the context of symbiotic or host parasitic interactions, but specificity can be more broadly used to describe the extent to which a species occupies a narrower range of an environmental variable than expected by chance. Using a standardization we describe here, Rao’s (1982, 2010) Quadratic Entropy can be conveniently applied to calculate specificity of a feature, such as a species, to many different environmental variables.

**Results:** We present our R package *specificity* for performing the above analyses, and apply it to four real-life microbial data sets to demonstrate its application. We found that many fungi within the leaves of native Hawaiian plants had strong specificity to rainfall and elevation, even though these variables showed minimal importance in a previous analysis of fungal beta-diversity. In Antarctic cryoconite holes, our tool revealed that many bacteria have specificity to co-occurring algal community composition. Similarly, in the human gut microbiome, many bacteria showed specificity to the composition of bile acids. Finally, our analysis of the Earth Microbiome Project data set showed that most bacteria show strong ontological specificity to sample type. Our software performed as expected on synthetic data as well.

**Conclusions:** *specificity* is well-suited to analysis of microbiome data, both in synthetic test cases, and across multiple environment types and experimental designs. The analysis and software we present here can reveal patterns in microbial taxa that may not be evident from a community-level perspective. These insights can also be visualized and interactively shared among researchers using *specificity*’s companion package, *specificity.shiny.*

## Introduction

The word “specificity” has uses across multiple disciplines. In ecology, and especially for microbes, “specificity” is often used in the context of symbiotic interactions; for example the specificity of a parasitic species may be the degree to which it associates with a narrow consortia of host species [1; 2; 3]. In pharmacology and biochemistry, specificity can describe the “narrowness of the range of substances with which an antibody or other agent acts” [4]. Synthesizing these definitions, we arrive at a general concept of specificity, where a feature *(e.g.* a species) is specific to some variable *(e.g.* elevation) when it occupies or is otherwise associated with a limited breadth of that variable.

This definition is consistent across multiple variable types. For example, a species that is found only across a narrow band of elevation, perhaps between 200 and 500 meters above sea level would have stronger specificity to elevation than a species that is found between sea level and 1000 meters. This is similar to a parasite that is found only within a narrow clade of hosts; it has stronger host specificity than a parasite that is found across a much wider phylogenetic range [2]. This concept can be expanded even farther, to diversity of some co-occurring feature class. For example, metabolites that co-ocur with bacteria in the human gut microbiome (microbes within the human gut). Under our definition, a microbe may have specificity to a narrow range of metabolomic compositions. Furthermore, specificity as we describe it here, is not the same as bipartite network specialization like *H*2′, *d*′, and *NODF* metrics [5; 6]. Those metrics apply to strictly categorical contingency data, for example a matrix of observation counts where columns are pollinator species and rows are plant species. Instead, our generalized specificity approach is best suited to continuous data.

Our generalized specificity analysis has several benefits over modeling a microbe’s relative abundance using a variable of interest, since specificity analysis has no underlying model. First, high-throughput sequencing (HTS) microbiome data notoriously contain many zeroes, corresponding to the lack of an observation of species in a samples. Disregarding the difficulties in modeling such data, which certainly can be overcome [7], these data are perfect for the “specificity approach”. This is because the alternative hypothesis of a specificity analysis (the focal species encounters less environmental heterogeneity than expected by chance) includes cases where the focal species only occupies a limited range of the variable of interest, being absent (zero) everywhere else. A further consideration in modeling approaches is non-monotonic relationships between species and environmental variables. For example, a species may have specificity to intermediate elevations, so its density function of elevation would be non-monotonic, or even multimodal; and that’s just one species. Within a HTS microbiome dataset, species may be expected to run the gamut of distribution shapes and modalities. Variables of interest also present their own challenges to modeling, since variables may be vectors *(e.g.* elevation, pH), distance or dissimilarity matrices *(e.g.* geographic distance, beta-diversity), phylogenies, or even sample-type ontologies. The generalized specificity approach we present here can accomodate all of the aforementioned variable types, unlike other approaches where the statistics used to understand microbe-environment relationships are restricted by variable type. Furthermore, our approach does not produce a model, or answer the question “across what range of the variable does the species occur”. Instead, we quantify the extent to which the species occupies a limited breadth of that variable without the need for such a model.

Meaningfully applying this general idea of specificity to multiple data types is challenging because of the different specificity metrics available to different kinds of data. With host phylogenetic data, specificity may be calculated as phylodiversity [8], or host phylogenetic entropy [9], or host richness [10]. However, with other data types these metrics are not useful – one cannot calculate phylogenetic entropy of elevation, for example. Per our definition above, specificity must be a measure of the breadth *(i.e.* heterogeneity, diversity) of an environmental variable occupied by the focal species. With a variable like elevation, a naive specificity metric may be as simple as the variance in elevation where the focal species is present, or weighted variance for a more intuitive approach. However, such a metric would not be applicable to phylogenetic data sets because it is limited to 1-dimensional data types (*i.e*. column vectors). Furthermore, we wanted our general idea of specificity to be useful for dissimilarity matrices. We found that Rao’s Quadratic Entropy [11; 12; 13] is a convenient diversity metric that can be applied to all abovementioned data, with a modicum of standardization (detailed in our Methods section).

Here, we present a software package written in R and C++ that implements a generalized specificity analysis. Our package, *specificity,* calculates specificity values for each species in a sample-by-species matrix. In microbiology, this data structure often appears as a table of OTUs (operational taxonomic units; a substitute for species) or ASVs (Amplicon Sequence Variants; OTUs represented by unique sequences after applying a denoiser such as DADA2 [14]). We simulated species distributions with varying strengths of specificity, and used those simulated data to validate our implementation. Our simulations were also used to ensure that specificity is not sensitive to occupancy (*i.e.* in how many samples a species appears), which is a significant improvement compared to the standardized effect size (SES) method [2; 15], and methods that use un-weighted (presence-absence) species data [10]. Our simulations also confirmed that the specificity we calculate here is scale-invariant with regard to environmental / phylogenetic data, and also to focal species abundance data. To illustrate how specificity can be used, we applied our software to four previously-published microbiological data sets, each from different environments: fungi living within the leaves of native Hawaiian plants, human gut microbiome bacteria, bacteria living within Antarctic glaciers, and the global Earth Microbiome Project data set.

## Methods

### RQE

We calculate specificity using Rao’s metric [11; 12]. It is sometimes abbreviated FDQ for quadratic functional diversity, but since we use the same mathematics in a non-functional context, here we simply refer to the metric as *RQE* (Rao’s Quadratic Entropy), similar to the use of “QE” by its inventor. *RQE* is the sum of the elementwise product of two square matrices (excluding the diagonal). In our use, the first matrix *(D*) is a dissimilarity matrix containing differences between samples (Figure 1). For example, in the context of phylogenetic specificity these differences are phylogenetic distances (*i.e.* cophenetic distances) between hosts. Samples from the same host species have 0 distance. The second matrix (*W*) contains all pairwise products of weights for the focal species. Given a column vector of species weights *p* from a site-by-species matrix (“OTU table”), *W_ij_* contains the product of the abundances (weights) of the focal species at sample i and sample j: *p_i_p_j_*. Via *D* o *W* (or *D_ij_p_i_p_j_* for a single pairwise product; Equation 1), we weight matrix *D* to up-weight distances between samples where the focal species occurred, and down-weight distances between samples where the focal species was absent in either. We use the term “weights” to describe *p* because the values within could be relative abundances or any other metric that describes the importance of a species within a sample. Conversely, we have chosen to focus this manuscript on “species”, but note that *p* could be a vector of weights for any feature (a type of rock, a metabolite, etc).

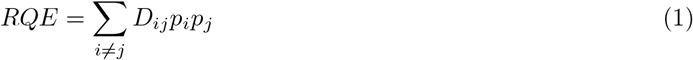

With *RQE*, a focal species with strong specificity has relatively high weights for low differences. This metric was originally developed for phylogenetic distances, but here we apply it to many different *D* matrices, including euclidean transformations of 1-dimensional data (e.g. pairwise elevational difference), or more complex 2-dimensional data like Bray-Curtis dissimilarity between host metabolomic profiles.

**Figure 1:**
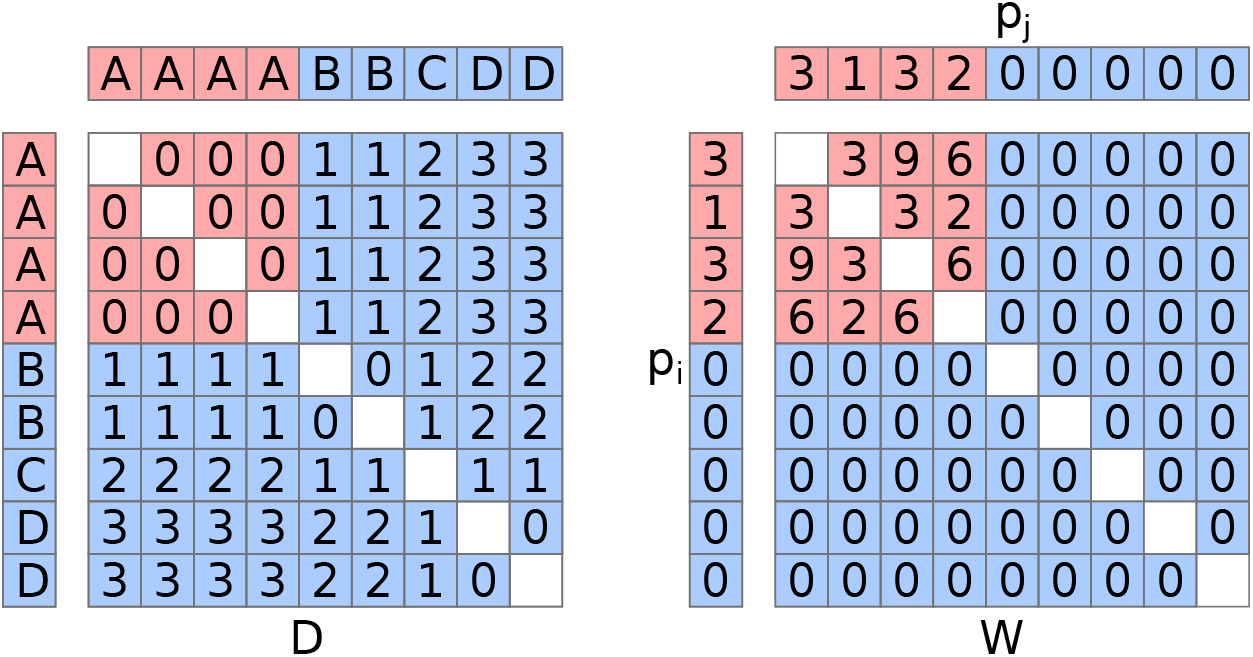
*RQE* as it applies to specificity. In this example, two matrices are shown, *D* and *W*. *D* is an environmental dissimilarity matrix, describing how different are several environment types, A through D, with multiple samples represented for each environment type. Note that diagonals are empty because they are not used; see Equation 1. Matrix *W* is the pairwise product of species weights *p* (Equation 1). In this example, the focal species is perfectly specific to habitat A, which can be seen in p. Data corresponding to species detections are colored in red, and species absences in blue. The product D o W (=*D_ij_p_i_p_j_*) will be all zeroes for this example, because this example shows perfect specificity. Thus, the sum of that product, *RQE*, will be zero. If *p* had relatively small values instead of its zeroes, for example 0.25, those small values would still down-weight their corresponding larger differences in *D* and produce a signal of specificity, compared to random permutations of *p* which produce much higher *RQE* values.

As such, a species with “perfect” specificity will always have *RQE* = 0. For example, consider a focal species *S* that can be found in habitats A, B, C, or D, with multiple samples collected for each category (Figure 1). If S is only found in samples from habitat A, matrices D and W will contain zeroes in opposite positions, resulting in *RQE* = 0. Note that weights near zero can also act similarly to zero since this is a weighted metric. In this way, a focal species can occupy every single sample (all values of *p* are nonzero positive) and still have *RQE* near zero. This is important so that spurious species detections do not significantly contribute to specificity. For example, in a DNA sequencing experiment, small amounts of contamination may occur during DNA extraction or library preparation. The magnitude of that contamination is expected to be small compared to the signal in an actual sample, but may result in spurious species detections. However, because these contaminants would be expected to be rare in the sample, their weights would be low in samples where they are noise, and high in samples where they are signal.

### Standardization

While empirical *RQE* is calculated as described above, it must be standardized in order to compare effect size between different variables and different species, because the metric’s scale is dependent on the scales of *D* and *p*. Phylogenetic specificity approaches have previously used a standardized effect size (SES) approach [2], but we found that SES has unfortunate properties when used with our generalized approach to specificity. Critically, SES was highly sensitive to occupancy, which is the number of samples a species occupies. One would expect the strength of specificity to be lower when a species occupies more samples, because this means the species must occur in a broader range of habitat. However, SES counter-productively yields stronger specificity for more occupant species, and also for species with more even distributions. This is because SES is standardized using a distribution of values generated with permuted species weights. If all weights are similar (high evenness), the standard deviation of that distribution will be small, leading to a strong SES. SES is also undesirable because for a given species, it is tightly associated with that species’ *P*-value (the probability of SES being as strong or stronger, despite the null hypothesis being true), enough so that a suggested remedy for yet other problems with SES is to use a probit transformed P-value in its place [15].

We standardize *RQE* by leveraging the fact that for perfect specificity, empirical *RQE (RQE_emp_,* Figure 1) equals zero. Our statistic, which we simply call *Spec,* ranges from −1 to 1, with 0 as the null hypothesis that species weights are randomly ordered with regard to sample identity. Similar to SES, *RQE_sim_* is a vector of *RQE* values calculated using random permutations of *p*. The distribution of *RQE_sim_* is used for calculating P-values, and its central tendency is defined as *Spec* = 0. This central tendency can be calculated several ways using our software, but the default is to use the mean of *RQE_sim_* since in our testing, mean and median showed highly concordant *Spec* results (Supplementary Figure S1). The equation for *Spec*(Equation 2) is a piecewise function, with the two parts corresponding to specificity and generality, respectively. In the case of specificity, *S_pec_* simply scales *RQE_emp_* relative to the center of *RQE_sim_* such that perfect specificity returns −1, and the null hypothesis returns 0. In this case, “null hypothesis” refers to *RQE_emp_* being the expected value of *RQE_sim_.*

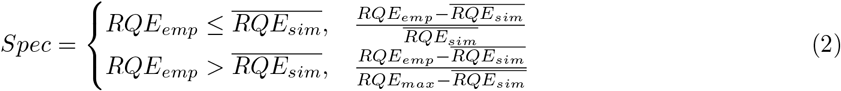

The case of generality is slightly more complicated, since there is no intuitive maximum theoretical*RQE*value. Generality in this context refers to species that encounter greater environmental heterogeneity than expected by chance. We find that maximum value computationally, and standardize *Spec* as a proportion of that value (see Equation 2). For each *p* there exists an optimized permutation that yields the highest possibleRQEvalue, *RQE_max_.* We use a genetic algorithm (GA) with Population Based Incremental Learning [16] to search permutations of *p* that create *RQE_max_.* Our GA begins with a population of surrogate vectors initialized via random permutations of *p* (default 150), and random swaps of *p* (a swap being the pairwise substitution of two values within the vector; default 150), and also *p* itself. Each generation, the GA calculates *RQE* for each vector in the population, then keeps some of the vectors with the highest *RQE* value (default 5). The next generation is composed of those kept vectors, and random swaps thereof until the total original population size is met (default 301). Our swapping algorithm can also use a stochastic number of swaps per vector per generation (including initialization), drawn at random from a user-defined set (default 1,1,2,3). In addition to swapping, mutation can be performed by crossover via the PMX algorithm [17], which is used because it incorporates both order and position of both parents, which is required for this problem. However, in our testing we found that crossover did not improve GA efficiency, so the default operation is not to perform PMX. The GA runs for a fixed number of generations (default 400), or until a number of generations have passed with no improvement (default 10). These parameters were chosen because they performed well on the data sets we analyze here, meaning that species reached the early termination condition.

Our GA is relatively computationally intensive, consuming the majority of computational time for a given specificity analysis even though it is only used for a minority of species. This is unlikely to be a concern on smaller data sets (i.e. a few hundred samples), but since many users may not be interested in “general” species, another option is to scale Spec for all species using the top half of Equation 2 instead. This is considerably faster, and the user can either discard “general” species as uninteresting, or choose to interpret *Spec* > 0 within an ordinal framework (a brief analysis showed the results of this approach and those of the GA are strongly correlated; Supplementary Figure S2).

### Hypothesis testing

For the Spec calculation above, a P-value may be calculated as the proportion of *RQE_sim_* values that are lower than *RQE_emp_.* The default operation of our software is to adjust *P*-values calculated for different species from the same variable for multiple hypothesis testing by applying the Benjamini-Hochberg procedure [18].

### Features of *Spec*

*Spec* captures signal of specificity to simulated vector, matrix, and phylogenetic data (Supplementary Figures S3, S4). It is insensitive to species occupancy (Supplementary Figures S5, S6) and is insensitive to the number of samples within a data set (Supplementary Figure S7). Spec is also scale-invariant, independently in regard to *p* and *D* inputs (Supplementary Figure S8). It is sensitive to multimodality, and multimodal species distributions are still detected as exhibiting significant specificity by *Spec* (Supplementary Figure S9).

### Validation analyses

Species were simulated with varying levels of specificity by drawing from a normal distribution centered on an artificial “optimum” environmental location (*e.g*. elevation of 300 meters). Varying specificities were achieved by widening the standard deviation of that distribution, or by mixing the normal distribution with varying proportions of a uniform distribution. Multimodal specificity was simulated similarly by combining multiple distributions. Specificity of simulated species was analyzed using our software. Occupancy of simulated species was increased or decreased by randomly substituting simulated weights with zero, and specificity was analyzed across an occupancy gradient using that approach. Real data (see Endophyte analysis, below) were randomly downsampled to test the sensitivity of *Spec* to sample size. Real data were also re-ordered to create simulated high-specificity species that use empirical distributions of weights, and then those simulated species were subjected to a swapping algorithm that gradually introduced entropy into the species. The swapping algorithm swaps values from two randomly selected positions in *p* (Equation 1). This was done recursively for 1000 generations (2 swaps per generation), saving *p* each time. Our software was then run on all vectors simulated this way.

### Analysis of endophyte data

Data from Hawaiian foliar endophytic fungi [19] were downloaded from FigShare. These are illumina MiSeq data of the Internal Transcribed Spacer (ITS) region of fungal ribosomal RNA, from 760 samples collected from the leaves of native Hawaiian plants across five islands in the Hawaiian archipelago. This data set is also included in our R package. The features under investigation in this analysis were fungal OTUs. Data were transformed (“closed”) using total sum scaling, and fungal OTUs present in fewer than 10 samples were excluded from specificity analysis, because low-occupancy data can be unreliable (Supplementary Figure S6). The remainder (416 OTUs) were run through our software using default settings except run in parallel using 20 CPU cores. Variables used in this analysis were NDVI (an index of vegetation density), elevation, evapotranspiration, rainfall, host plant phylogeny, and geographic distance between sample sites.

### Analysis of Antarctic bacteria data

Data from Antarcic cryoconite hole bacteria [20] were obtained from the authors. Cryoconite holes are isolated melt pools on the surface of glaciers, caused by debris from nearby slopes falling onto the glacier, and then melting into its surface. These holes form discrete microbial communities that have been described as “natural microcosms”[21]. This data set comprises 90 samples across three adjacent glaciers, and features are bacterial (16S rRNA) and eukaryal (18S rRNA) Amplicon Sequence Variants (ASVs; a type of OTU). Taxonomy was assigned to 18S rRNA ASVs using dada2 [14], and Bray-Curtis beta-diversity was calculated for only those ASVs that were determined to be algae. Analyses on 16S rRNA data were run and visualized as above, with variables N (Nitrogen), P (Phosphorus), pH, geographic distance, fungal Bray-Curtis beta-diversity (calculated from 18s rDNA data), and algal beta-diversity. Bacterial associations with individual glaciers (*e.g*. “OTU4 is found predominantly on Canada Glacier”) were computed using Dunn’s test [22], which is a nonparametric post-hoc test of difference in means.

### Analysis of Human microbiome data

Data from Franzosa et al. [23] were downloaded as supplemental data from the online version of the article. These data contain both gut bacterial and archaeal species composition data as well as corresponding metabolomic data, collected from 220 adults with Crohn’s disease, ulcerative coliutis, or healthy controls. Data were downloaded in a processed state, after the following procedures had been completed: species composition data from this study were derived from metagenomic data, which were assigned taxonomy and grouped into OTUs using MetaPhlAn2 [24], and excluded samples that did not meet a 0.1% relative abundance threshold in at least 5 samples. Metabolite data were measured using positive and negative ion mode LC/MS, and were reported as parts per million. Metabolite identities were assigned programatically, and were clustered into broad classes per the Human Metabolome Database [25]. We subset the matrix of metabolome data by those classes, and used Euclidean distance to calculate the extent to which any two given samples differed in metabolomic composition within a given class (*e.g.* “how different is the composition of bile acids between sample A and sample B?”). Metabolite classes were excluded if they were totally absent in any sample, or if they contained fewer than 10 metabolites, which left 83 classes. specificity was used to calculate *Spec* for microbial OTUs to each metabolite class distance matrix.

### Analysis of Earth Microbiome Project data

Data from the Earth Microbiome Project (EMP)[26] were compiled and downloaded from Qiita [27]. These data comprise a global sampling of 16S rRNA ASVs produced by multiple studies. All of the studies followed a uniform protocol for collection, processing, and analysis of microbial data. A major component of the EMP is a rigid sample type ontology. The EMP Ontology (EMPO) was designed to categorically represent two main drivers of bacterial community composition: host association and salinity, for each sample that was collected. At the broadest level (EMPO1), samples were categorized as being host-associated or free living. At the intermediary level (EMPO2), samples were further divided into saline, non-saline, animal, plant, and fungus. The finest level (EMPO3) separated samples into 22 discrete substrate types (e.g. water (saline), plant corpus, animal distal gut).

Because this data set is so large (28,842 samples and 309,469 ASVs), ASVs were excluded that were not present within at least 30 samples, samples were discarded if they had fewer than 5,000 reads, and samples without ontological data were discarded (leaving 25,188 samples and 7,014 ASVs). ASVs were excluded due to low occupancy to avoid spurious ASVs and to avoid low-abundance ASVs that do not perform well with *Spec*, and more importantly to keep computation size manageable for this massive data set. Samples were also discarded mainly due to computational concerns, with low-count samples being dropped first due to lower confidence in their proportional abundance calculations. The EMP ontology was transformed into a phylogeny using specificity’s “onto2nwk” function, which makes a cladogram within which all branch-lengths were set to 1. Specificity analysis was run using the ASV table and the ontological data. Database matches for individual species of interest were manually obtained using nucleotide BLAST [28] via the NCBI web portal, using the 16S rRNA sequence database as reference.

### Implementation

*specificity* was written in the R programming language, with some functions written in C++ using *Rcpp* [29]. The general format of the package follows standard R package structure [30]. Unit testing was done using *testthat* [31]. *specificity.shiny* was written entirely in R, and uses the *shiny* [32] interactive web application framework. Both packages are free and open source software, licensed under the Gnu Public License (GPL). Installation is easily done using the “install_github” function of the R package remotes [33]; see data availability section for details.

## Results and discussion

### Hawaiian endophyte specificity analysis

We found that foliar endophytic fungi (FEF) from within the leaves of native Hawaiian plants exhibited strong and statistically significant specificity to several environmental variables (Figure 2), including variables that were only weakly associated with FEF community composition [19]. For example, in the original paper, rainfall and elevation were relatively weak predictors of FEF phylogenetic beta-diversity, but many FEF species show strong specificity to those variables in the analysis presented here. This reflects a fundamental difference between community-centric approaches (*e.g*. FEF community composition) vs. species-centric approaches like (*e.g.* specificity analysis). The signal of individual species is lost when a community is aggregated into a beta-diversity matrix or similar, and consequentially individual species within the community may even respond to environmental variables orthogonally to the community as a whole. Species that were strongly specific to rainfall or elevation are examples of this.

**Figure 2:**
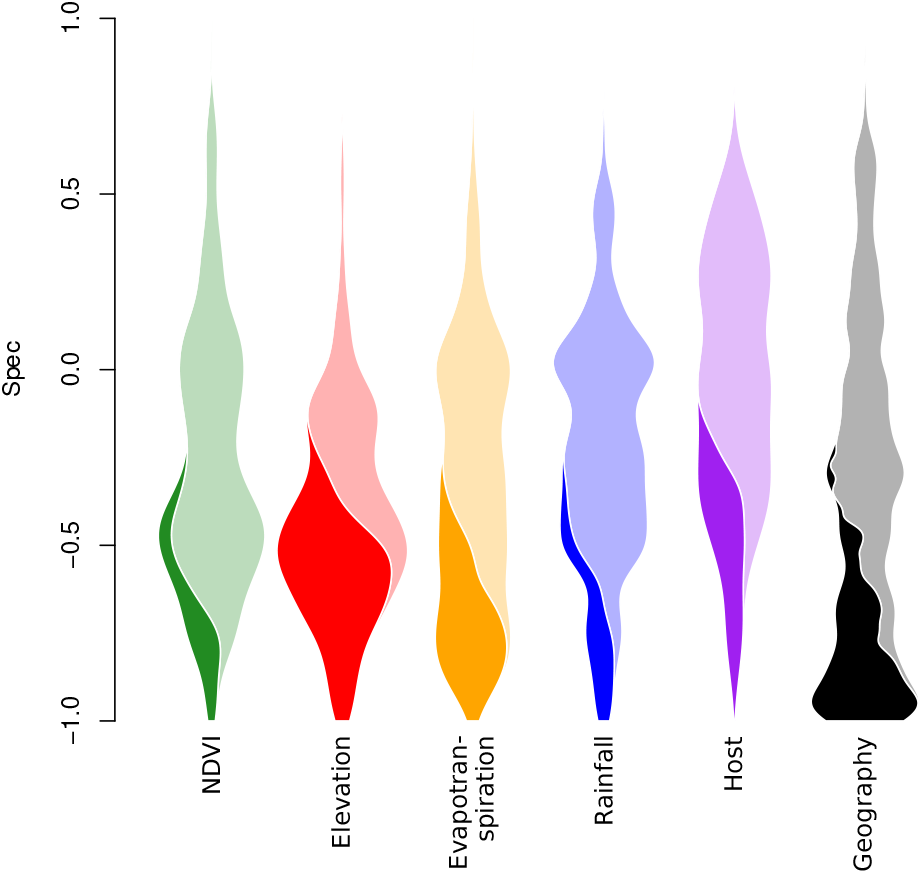
Specificity of Hawaiian Foliar Endophytic Fungi. In this plot, the *Spec* values for 416 fungal species are plotted as violins for different variables. Since the number of species is the same for each variable, each violin has the same total area. Violin area is divided between species with statistically significant specificity (dark) vs. species without (light).

We found that many FEF species have strong and statistically significant specificity to geographic location, which makes sense given the discrete spatial structure of the Hawaiian islands [34], and that these FEF communities only are spatially structured up to distances of 36 kilometers [19]. But geographic specificity may be an artifact of specificity to other variables with strong geographic autocorrelation. For example, in Hawaii (and elsewhere), rainfall is a spatially structured phenomenon [35], with nearby areas experiencing dramatically different rainfall averages as a consequence of aspect and elevation. Thus, species that have strong specificity to rainfall likely also have strong specificity to geography, which is true in our analysis. Indeed, the same is true for elevation, albeit to a lesser extent Figure 2.

One of the fungi with strongest specificity in our analysis was of the genus *Harknesia,* with closest BLAST match in the NCBI nucleotide database to *H. platyphyllae,* a eucalyptus pathogen. In our data set, this fungus was found on multiple hosts, including *Metrosideros polymorpha,* which is in the same family as eucalyptus *(Myrtaceae*). This *Harknesia* was found exclusively within the interior of Hawaii island, at elevations between 1700 and 2000 meters above sea level. Likely because of its strong geographic and elevational specificity, it also exhibited strong specificity to evapotranspiration, only being found in areas with 40 to 50 mm per year. Other fungi, such as an ASV most closely related to *Phaesosphaeria papayae,* exhibited strong specificity to elevation, evapotranspiration, and vegetation (NDVI), but were found on multiple islands across the Hawaiian archipelago. Thus, while geographic specificity can appear as specificity to geographically autocorrelated variables, this is not always the case.

Notable generalists *(Spec* > 0) in the endophyte dataset include the genus *Colletotrichum,* a globally distributed genus of plant pathogenic and endophytic fungi. Almost all agricultural crops are impacted by members of *Colletotrichum* and it is considered a ‘top ten’ fungal pathogen for molecular plant pathological research [36]. Of the 9 ASVs identified as *Colletotrichum,* none showed specialization to plant host or geography. Recently, genomic studies of this genus have provided insight into the genetic mechanisms behind host generalism and the activation of latent pathogenicity [37; 38]. Low specificity to geography and host within this dataset indicates that asymptomatic *Colletotrichum* species are widespread within the native Hawaiian flora.

### Antarctic glacier bacteria specificity analysis

Similar to the FEF analysis above, bacteria living in cryoconite holes (isolated melt pools) on glaciers in Antarctica’s Taylor Valley [20; 21] exhibited strongest specificity to geographic distance (Figure 3). This data set spanned three glaciers: Canada, Taylor, and Commonwealth, with equal sampling on each, but geographic distance accounts for distances within glaciers as well. The strong geographic specificity observed here reflects bacteria that are differentially abundant among glaciers, for example occupying only one or two of the three. The three glaciers, each flowing into a separate lake basin, are spaced along the 40-km length of the Taylor Valley, which stretches from the polar ice sheet to the coast. The cryoconite holes from the most inland glacier, Taylor Glacier, have fewer nutrients than those nearer the coast, on Commonwealth Glacier [39]. The more inland cryoconite holes also have the lowest diversity of bacteria, while the holes nearest the coast support the most diverse bacterial communities [21]. Many bacterial species may therefore be specific to Commonwealth Glacier because it supports more species within its cryoconite holes than the other two glaciers. Besides the differences in bacterial richness among the glaciers, the composition of the bacterial community turns over among glaciers, with different sequence variants dominating each glacier [20]. Biogeochemical differences within cryoconite holes among glaciers furthermore correspond with biogeochemical gradients along the valley in the surrounding soils [39]. The difference in dominant bacterial taxa on each glacier may primarily reflect (1) differences in which bacteria dominate the soils surrounding each glacier and therefore disperse onto each glacier, (2) a response to biogeochemical conditions within cryoconite holes on the glacier, or (3) an interaction of the two. Experimental microcosms manipulating dispersal and nutrient availability could help to parse out dominant controls on geographic specificity of bacteria in the future.

**Figure 3:**
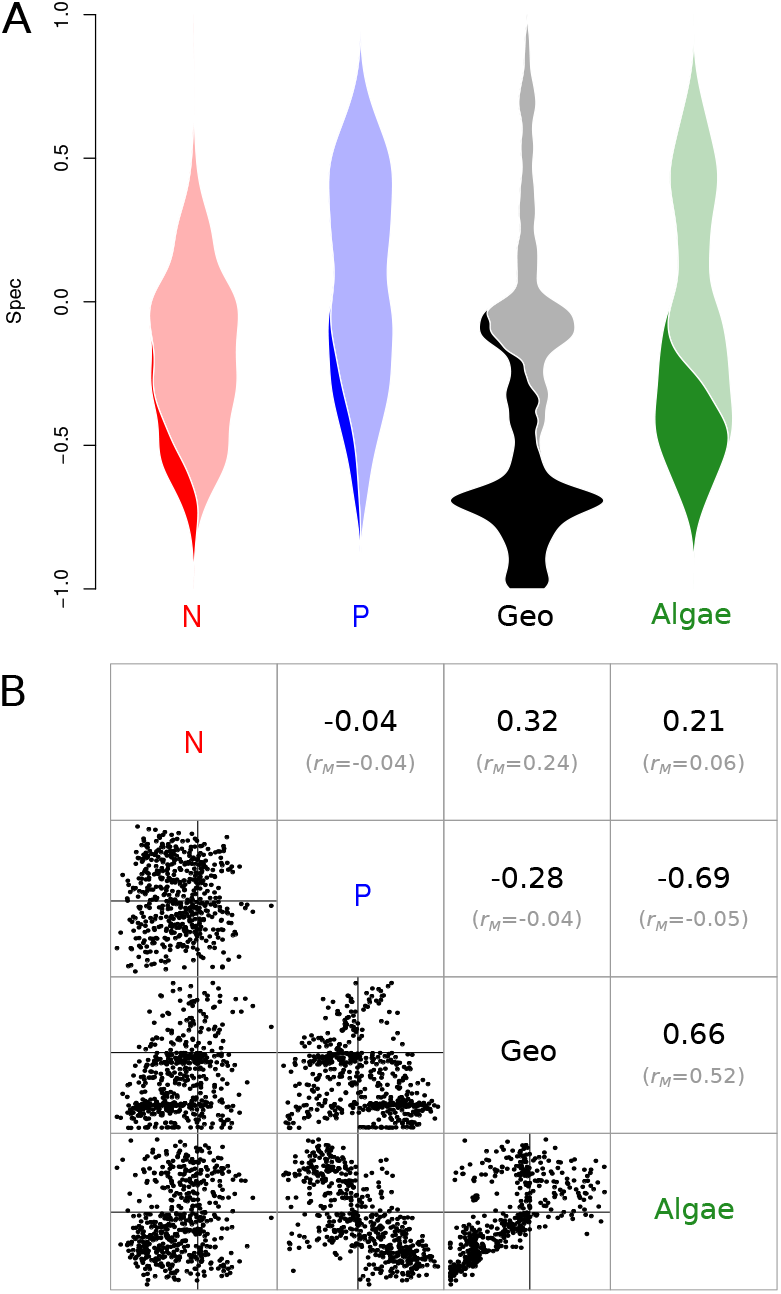
Specificity of Antarctic cryoconite hole bacteria. **A**: Violin plot of specificity to sediment nitrogen concentration (N, red), sediment phosphorus concentration (P, blue), geographic distance (Geo, black), and algal community composition (Algae, green). Violins are scaled and shaded identically to those in Figure 2. **B**: Pairwise *Spec* correlations. Correlation coefficients (r) for each pairwise comparison are shown in this subplot’s upper triangle, with mantel correlation coefficients *(r_M_*) shown below in gray. Mantel correlations describe the relationship between the variables themselves, *e.g.*“is algal community dissimilarity correlated with geographic distance”. The Spec correlations shown above in black are the correlation values for the data plotted in the lower triangle of this subplot.

Strong bacterial specificity to co-occuring algal communities was expected, given the strong correlation between bacterial and eukaryal diversity previously observed in this supraglacial system [20] and elsewhere [40]. In our analysis, we found that specificity to algae was strongly negatively correlated with specificity to phosphorus (*r* = −0.69); even though those two variables were not strongly correlated with each other (*r_M_* = −0.05). In other words, bacteria that are specific to algal community composition are not specific to sediment phosphorus concentration, and vice-versa. Using post-hoc tests, we found that bacteria with strong specificity to phosphorus concentration were predominantly associated with Taylor Glacier (but not exclusively), and that bacteria with strong specificity to algal community composition were predominantly found on Commonwealth Glacier.

Similarly, the correlation in *Spec* between geographic distance and algae (Figure 3B) highlights a feature of specificity analysis using *Spec*: when comparing specificity of the same features to two different variables, *Spec* is likely to be strongly correlated when the variables have a linear relationship. That linear relationship can be seen in the Mantel correlation between pairwise geographic distance and algal beta-diversity (rM=0.52; Figure 3B). But with variables that are weakly correlated, Spec may or may not be correlated between the variables. For example, difference in phosphorus concentration is not correlated with algal beta-diversity, but bacterial specificity to those variables is correlated (Figure 3B).

### Human gut microbiome metabolomic composition specificity

In this analysis, we asked whether bacteria and archaea in the human gut microbiome had specificity to paired metabolomic data. We computed bacterial specificity to compositional dissimilarity of 83 different metabolomic classes, and of those 83, microbes showed statistically significant specificity to only 25 (Figure 4). The interpretation of this analysis is similar to that of the specificity to algal community composition above. Specificity to a certain composition of paired data is a more abstract concept than specificity to elevation or even to the EMP ontology, but this type of specificity makes intuitive ecological sense in the context of species-habitat association. Different microbes have different environmental needs, both within the human gut microbiome [41] and elsewhere [42]. As such, those microbes can be expected to be found in environments that meet those needs. Similarly, microbes in the human gut influence their environment [43], and as such can be expected to be found in environments that are changed by their presence. Since different microbes interact with different sorts of metabolites, differential specificity to metabolite classes is an expected outcome.

**Figure 4:**
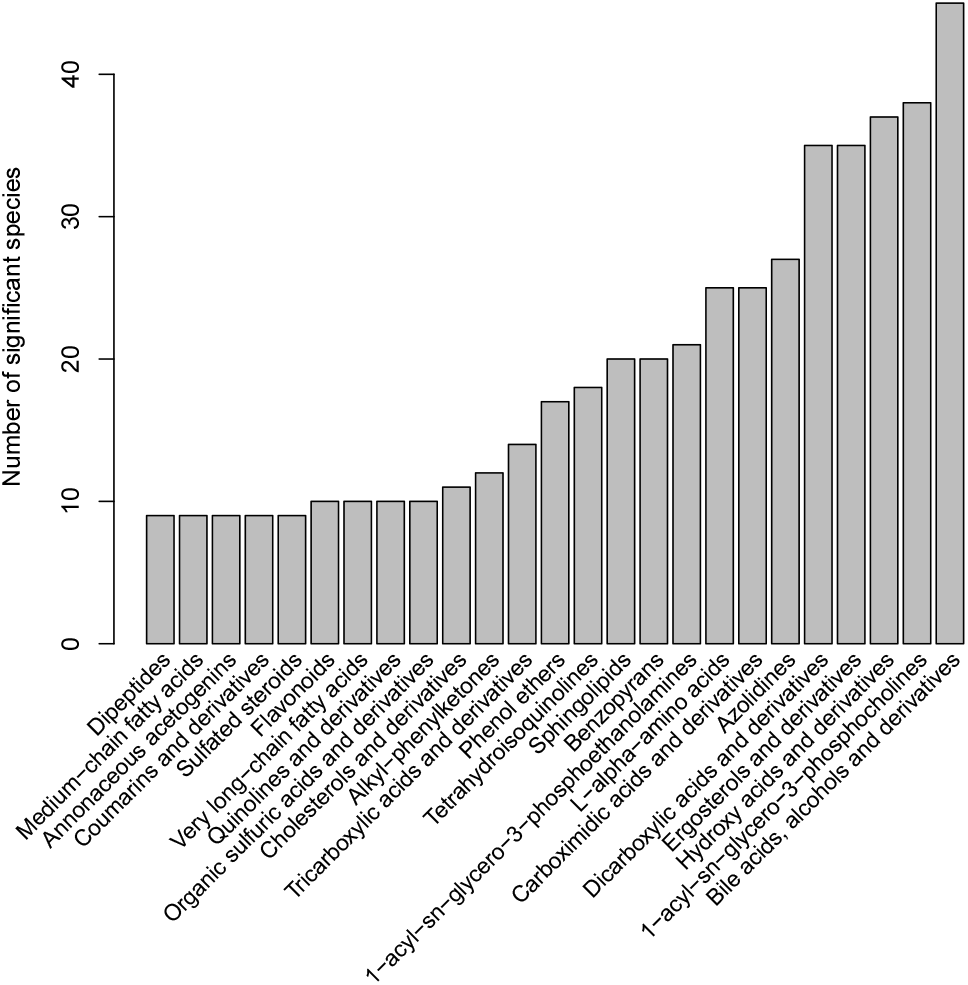
Count of bacterial and archaeal species that exhibited significant specificity to metabolomic composition in human gut microbiome data. Each bar represents the number of species (out of 201 total) that showed significant specificity to the composition of a given metabolomic class. Ordering of metabolomic classes on the horizontal axis is arbitrary.

We found that more microbial species had significant specificity to the composition of bile acids, alcohols, and derivatives, compared to other metabolomic classes (Figure 4). This result is not surprising, since bile acids strongly interact with the gut microbiome, and are also created and manipulated by it [44]. Furthermore, the experimental design for these data contained subjects with Crohn’s disease, with ulcerative coliutis, and healthy controls; and bile acids play a significant role in both Crohn’s disease and ulcerative colitis [45]. Microbes in this analysis could be specific to either of those two conditions and their plausibly co-occurring bile acids, alcohols, and derivatives, or to subclasses thereof. Composition of this metabolomic class did not strongly correlate with composition of other metabolomic classes; its highest Mantel correlation was with Benzopyrans (*r_M_* = 0.46).

Species with the strongest specificity to bile acids *etc.* were *Bacteroides plebeius,* an unclassified *Methanobrevibacter* species, *Odoribacter laneus, Methanosphaera stadtmanae,* and *Ruminococcus callidus* (all *Spec* < −0.60), although many other species showed significant specificity to this metabolomic class as well. *B. plebeius* was initially isolated on bile media [46], and was found to be associated with primary bile acids (as opposed to secondary bile acids) in patients with pediatric Crohn’s disease [47]. *Methanobrevibacter sp.* (likely *M. smithii*) and *Methanosphaera stadtmanae* (both Archaea) are the predominant methanogens found in the human gut [48]. *M. smithii* is known to grow in the presence of bile salts [49], and may be a biomarker against inflammatory bowel disease (IBD) or for its remission [50]. The metabolomic class with the second most specific microbes was 1-acyl-sn-glycero-3-phosphocholines (also called 2-lysolecithins). These compounds are derivatives of phosphatidylcholine, which is used as a treatment for ulcerative colitis, but is also found naturally in some foods [51]. This finding is different than showing some bacterial species’ relative abundances correspond to the amount of phosphatidylcholine derivatives; instead this analysis focuses on the composition of phosphatidylcholine derivatives; albeit with the amount of those compounds as a component since Euclidean distance was used.

In our analysis, we asked which metabolomic classes had the most microbial species specific to them (Figure 4). The results of this type of analysis are intended to mirror common beta-diversity analyses used in microbial ecology, which ask to what extent variables explain differences in microbiome community structure [19; 20; 52]. However, more complex questions can be asked of these data, using the results of specificity as a starting point for feature set reduction or variable selection. For example, given an individual bacterial species of interest, the variables to which it is specific may be used in a random forests model to predict its presence. For the purpose of variable selection, specificity has very low computational resource requirements when used with only the top half of Equation 2 (using option denom_type=“sim_center”), and can be run on personal computer hardware. This mode produces the same P-values as the more comprehensive mode, and produces the same *Spec* values for any species with *Spec* < 0. In addition to variable selection, specificity has application in detecting species that may be common lab contaminants, as shown in our EMP analysis below.

### Earth Microbiome Project (EMP) ontological specificity

As expected, the vast majority (6, 909/7,014) of bacterial ASVs we analyzed within the EMP data set exhibited significant and strong specificity to the EMP ontology (Figure 5). Given the distinct community-level differences in microbiomes across this ontology [26], it is not a surprise that most microbial species exhibit the same pattern. Instead, the species that buck this trend are of interest as potential cosmopolitan taxa, or possible bioinformatic failures. The sequence data obtained from the EMP data set are only 91 base-pairs in length, and even though they were clustered as exact sequence variants [14], it is possible that environmentally divergent ecotypes [53] were clustered together into the same ASV in this way.

**Figure 5:**
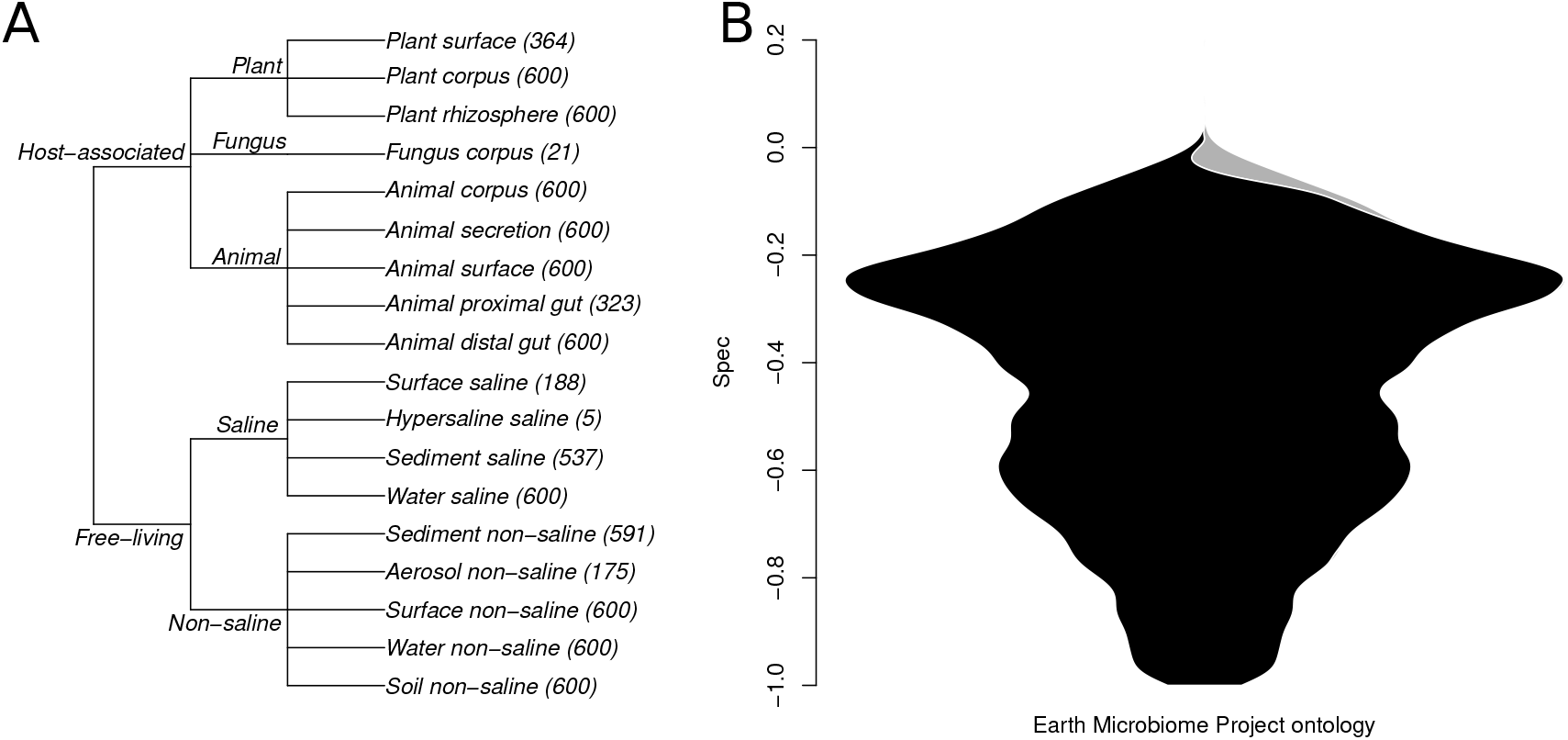
**A**: EMP sample type ontology visualized as a cladogram. **B**: Distribution of *Spec* values for ontological specificity of bacteria and archaea in the EMP data set. Within the cladogram, all brach-lengths are set to 1 for calculation of a cophenetic distance matrix, which is used as *D* when calculating *Spec* (Equations 1 and 2). Numbers in parentheses are sample counts. An overwhelming majority of microbes within the EMP data set showed statistically significant specificity to sample type ontology (black), which was expected given the previously reported strong signal in clustering of community composition by sample type [26]. Very few microbes did not show significant specificity (gray).

One such highly distributed ASV was found in almost all EMP ontology categories except for saline, hypersaline, and non-saline water (*Spec* = −0.13, *P* = 0.35). It was 100% identical to multiple *Actinomadura*species, including *A. algeriensis* from Saharan soil [54], the human pathogen *A. madurae*[55], the root endophyte *A. syzygii*[56], *A. maheshkhaliensis* from mangrove rhizosphere soil [57], and *A. apis* from honeybees [58], and others. These environment types span the EMP ontology, suggesting that the highly distributed ASV in question is a spurious combination of multiple *Actinomadura* ecotypes, each of which would likely have a strong specificity signal if analyzed independently. A counterexample is a strongly specific ASV, found exclusively in animal distal guts. It was 100% identical to multiple species as well, although each was originally isolated from similar environments: *Oxobacter pfennigii* (cattle rumen [59]), *Proteiniclasticum ruminis* (yak rumen [60]), and *Lutispora thermophila* (solid waste bioreactor [61]). These examples illustrate that like with every analysis, results can only be as good as input data. Users with very short-read marker gene data are likely already aware of this limitation, so we will not belabor the point.

### Interactive data visualization

In addition to using our *specificity* R package to calculate *Spec* and produce the figures shown above, we also used its companion package, *specificity.shiny*, to explore data and identify interesting features (Supplementary Figure S10). With this tool, users can easily create interactive visualizations from specificity analyses, and share them over the internet. *specificity.shiny* was used in the preparation of this manuscript, to share results between authors.

## Conclusion

Our R package, *specificity*, enables specificity analysis of microbiome data in the context of multiple variable types. Here, we’ve shown examples of specificity to geographic variables like elevation and rainfall (Figure 2), host phylogenetic specificity (Figure 2), specificity to co-occurring microbial community structure (Figure 3) and metabolomic structure (Figure 4), and specificity to sample type ontology (Figure 5). Our validation analyses show that our statistic, *Spec*, performs intuitively and is sensitive to specificity in both empirical and simulated data (Figures S3-S9). Our companion package, *specificity.shiny,* can be used to explore results and collaborate on specificity analyses (Figure S10), which was done by the authors on the four example analyses we presented here. Both *specificity* and *specificity.shiny* are available from the authors’ GitHub repository, along with installation instructions and a tutorial vignette.

## Declarations

### Ethical Approval and Consent to participate

N/A

### Consent for publication

All authors have given consent for this manuscript to be published.

### Availability of data and materials

R packages *specificity* and *specificity.shiny* can be downloaded from GitHub: https://github.com/darcyj/specificity, https://github.com/darcyj/specificity.shiny. In addition to software, our *specificity*’s GitHub repository contains thorough documentation, including guidance on installation, and a full tutorial vignette for using specificity with examples from an included data set. Code and data to replicate the analyses shown here can be found on GitHub as well: https://github.com/darcyj/specificity_analyses.

### Authors’ contributions

JD wrote the main manuscript, with contributions from AA, SS, PS, and CL. Data analysis, figure preparation, and software development were done by JD. All authors reviewed the manuscript.

## Acknowledgements

The authors thank J. Siebert and C. Martin for many helpful discussions and data wrangling.

## Authors’ information

N/A

