## supplementary for "*specificity*: an R package for analysis of feature specificity to environmental and higher dimensional variables, applied to microbiome species data"

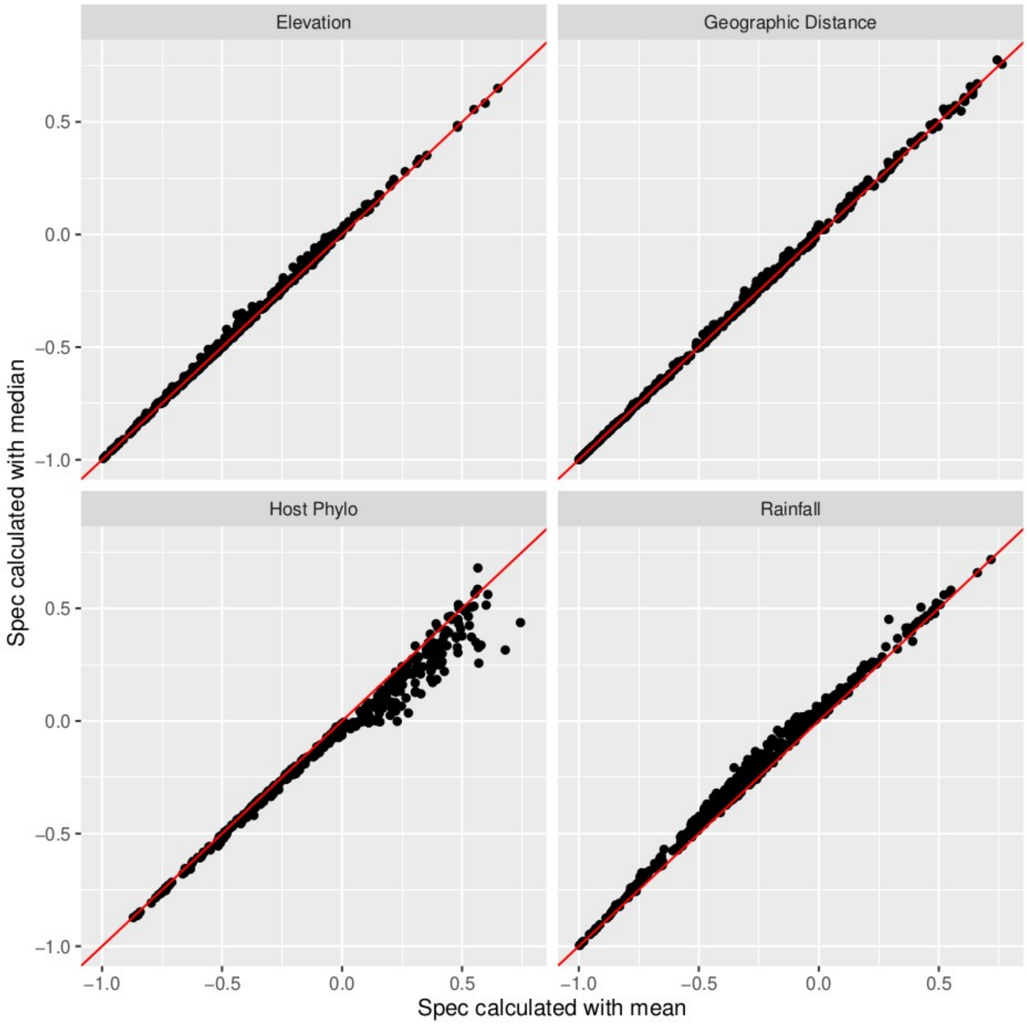


Supplementary Figure S1: Concordance of *Spec* when calculated with mean vs median as the measure of central tendency. In the cases we show here from our Endophyte data set, *Spec* shows strong concordance between runs using mean v.s. median. Red lines are 1:1 lines.

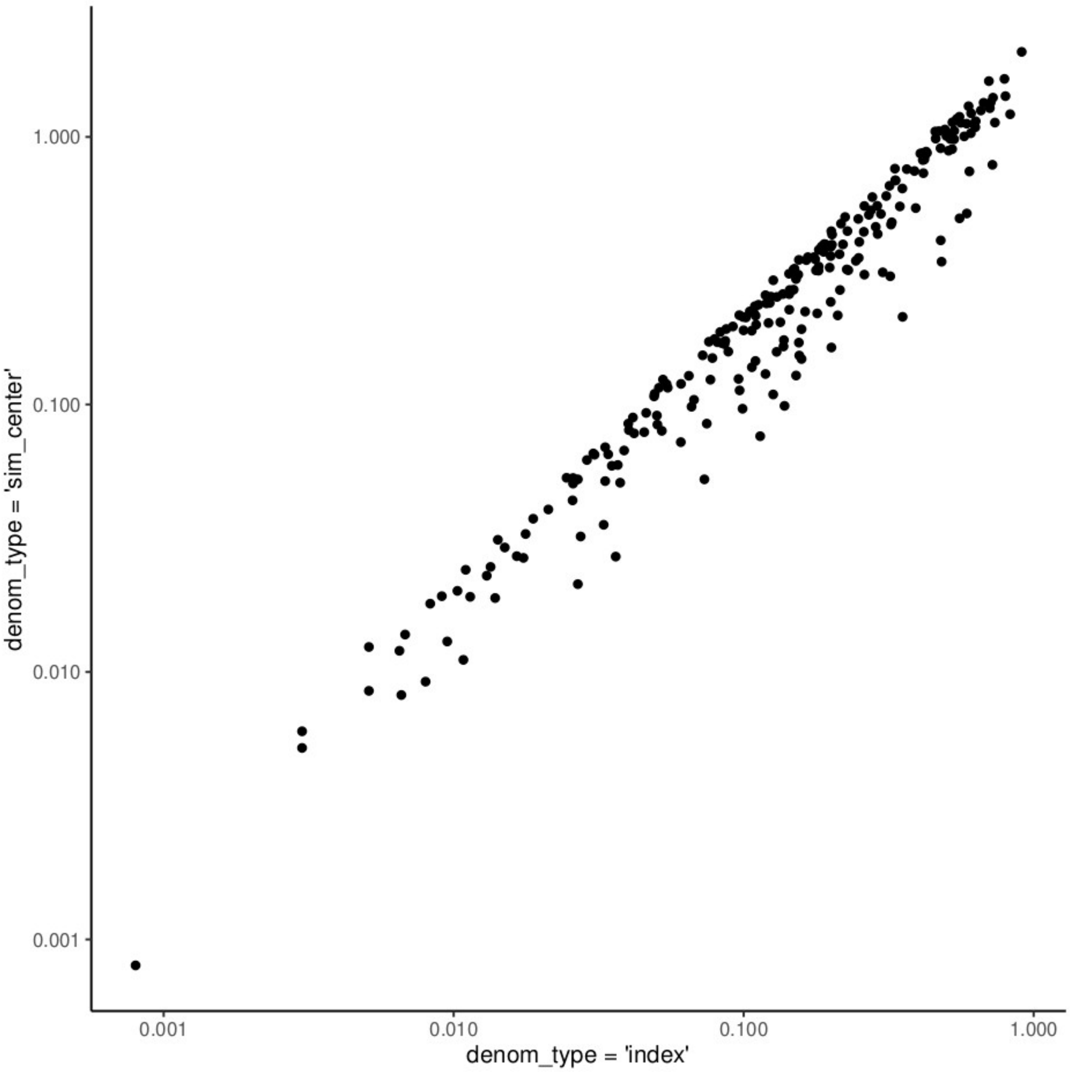


Supplementary Figure S2: Correlation of genetic algorithm (GA) and unscaled results for “general” species. In *specificity*, “general” features can be re-scaled using the GA approach, although this is computationally intensive. Users who are not interested in scaling “general” features can instead opt to scale all features using the top half of Equation 2, which is considerably faster. Results of both approaches are plotted above, with the GA approach on the horizontal axis, and the fast approach on the vertical axis. Note that both axes are log10 scaled to better visualize points; the actual scales of the two axes differ by a factor of two in this case, but would differ by different factors depending on what variable is used.

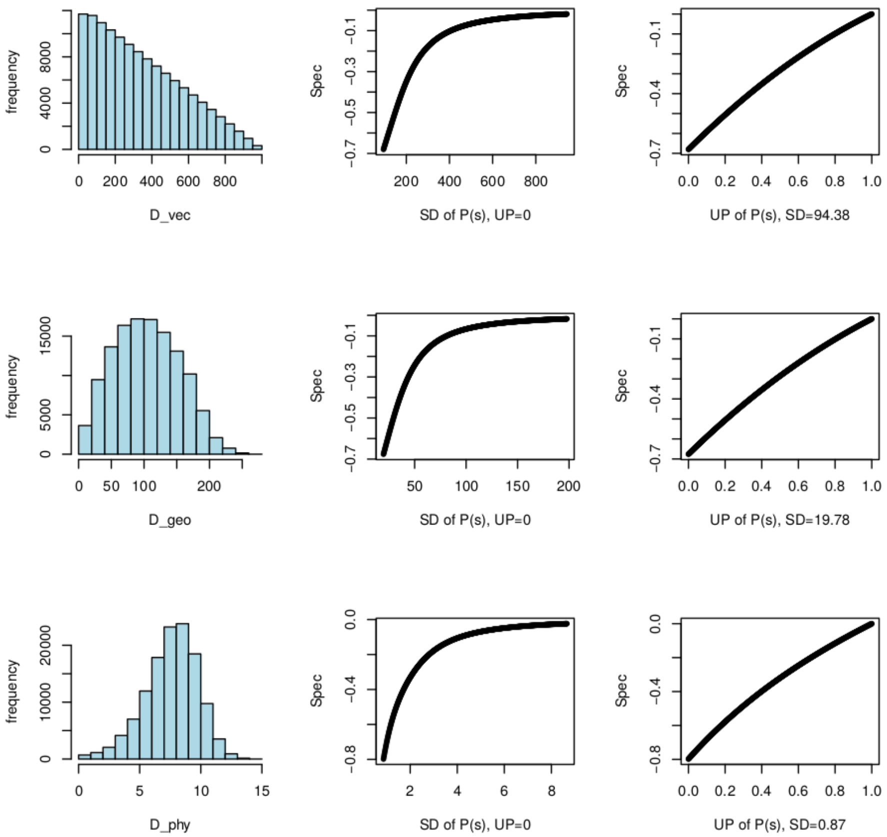


Supplementary Figure S3: Sensitivity of *Spec* to simulated specificity in vector, geographic (matrix), and phylogenetic data. Environmental variables were simulated as vector input (integers from 0 to 1000), geographic input (a random grid uniformly distributed points between -100 and 100 on both axes), and phylogenetic input (a random phylogenetic tree with 200 tips, generated using the *ape* R package, with tips sampled evenly). *D* (Figure 1) matrices for vector and geographic input were constructed using Euclidean distance, and cophenetic distance was used for phylogenetic input. An "optimum" sample was chosen at random. Species weights were simulated by constructing a probability distribution over each sample's distance (in *D*) to the optimum, using an additive mixture of normal and uniform distributions. The relative density contributions of the two distributions to the mixture was weighted by the parameter UP ("Uniform Proportion"); a distribution with UP=1 gives a uniform distribution, and UP=0 gives a normal distribution. The distribution's mean was fixed at zero (zero distance to optimum), then either the SD was varied to make the distribution narrower/wider, or UP was varied to make the distribution curvier/flatter. Strong specificity is represented by a distribution with low UP and low SD, but both were varied in order to test whether a focal species could have a background presence in all samples (given by the uniform distribution) and still exhibit specificity due to higher weights in a narrow subset of samples. SD values ranged between 0.4*SDD and 4*SDD, where SDD is the standard deviation of the lower triangle of *D*. UP values ranged between 0 and 1. For analysis varying UP, SD was held constant at 0.4*SDD. Species weights (*p*) were simulated as probabilities given *D*, and *Spec* was calculated from the simulated data. Results show that *Spec* is sensitive to unimodal specificity in all three data types analyzed, both by widening the distribution via SD, or attenuating the distribution via UP. Histograms for *D* are shown in the left column, sensitivity of *Spec* to SD in the middle column, and sensitivity of *Spec* to UP in the right column.

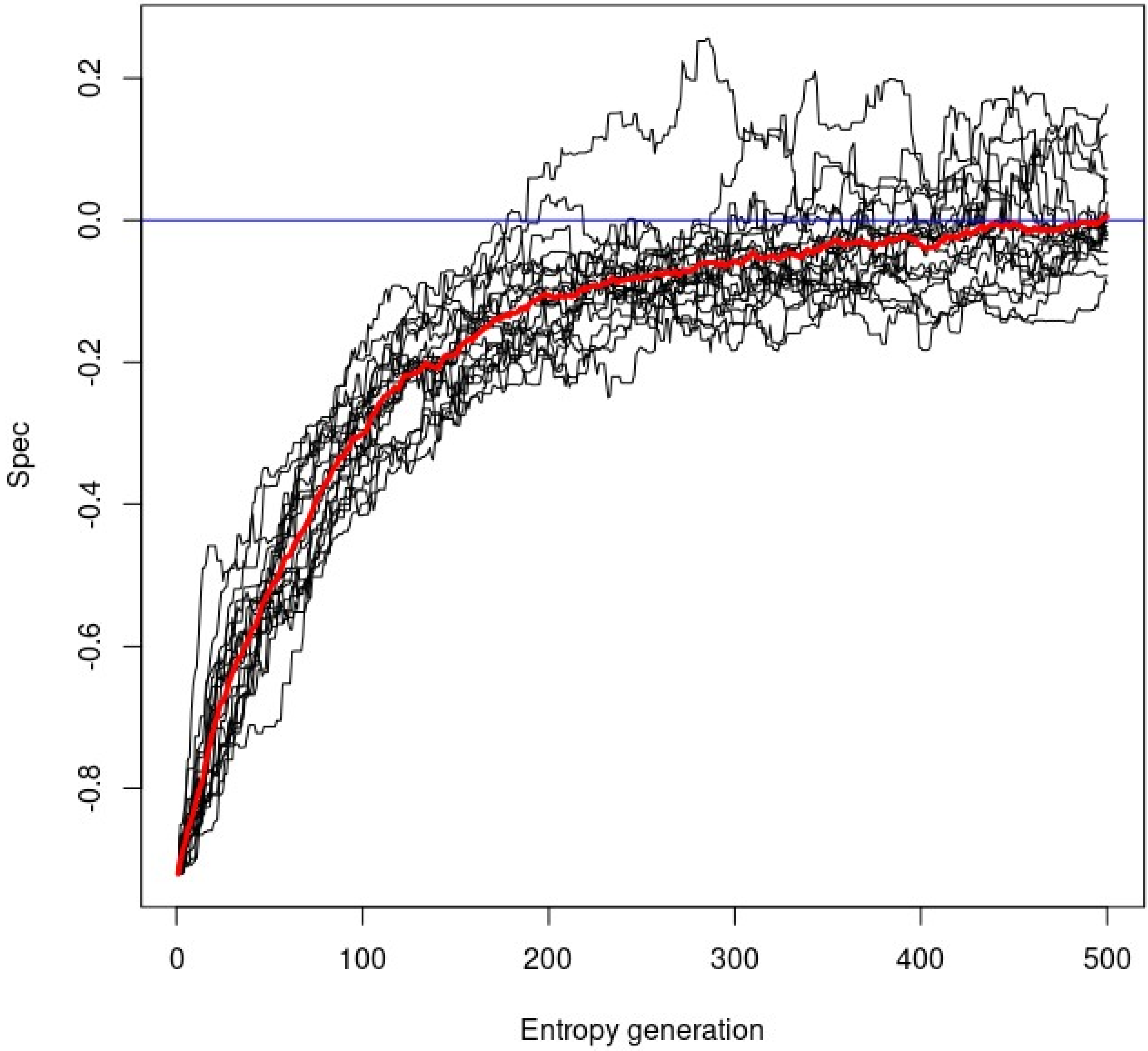


Supplementary Figure S4: Sensitivity of *Spec* to specificity as assayed via random perturbation of *p* (Equation 1). In this analysis, an empirical *p* vector was first sorted low-to-high to create a strong signal of specificity. Then, entropy was added to that vector by recursively swapping the position of two values within the vector. Three swaps were conducted per generation (x-axis), and *Spec* was calculated on the resulting *p*, using the Elevation variable from the Endophyte data set as the environmental variable. This procedure was run 20 times; each is visualized as a connected trend in the figure. If *Spec* is sensitive to specificity, moving from a highly specific *p* toward a uniformly randomized *p* will cause *Spec* to move from a negative value toward zero, then saturate as randomization reaches an equilibrium. This is what is shown in the figure, for a single fungal OTU. We ran this same analysis for several other OTUs and observed the same result. The red line is the mean.

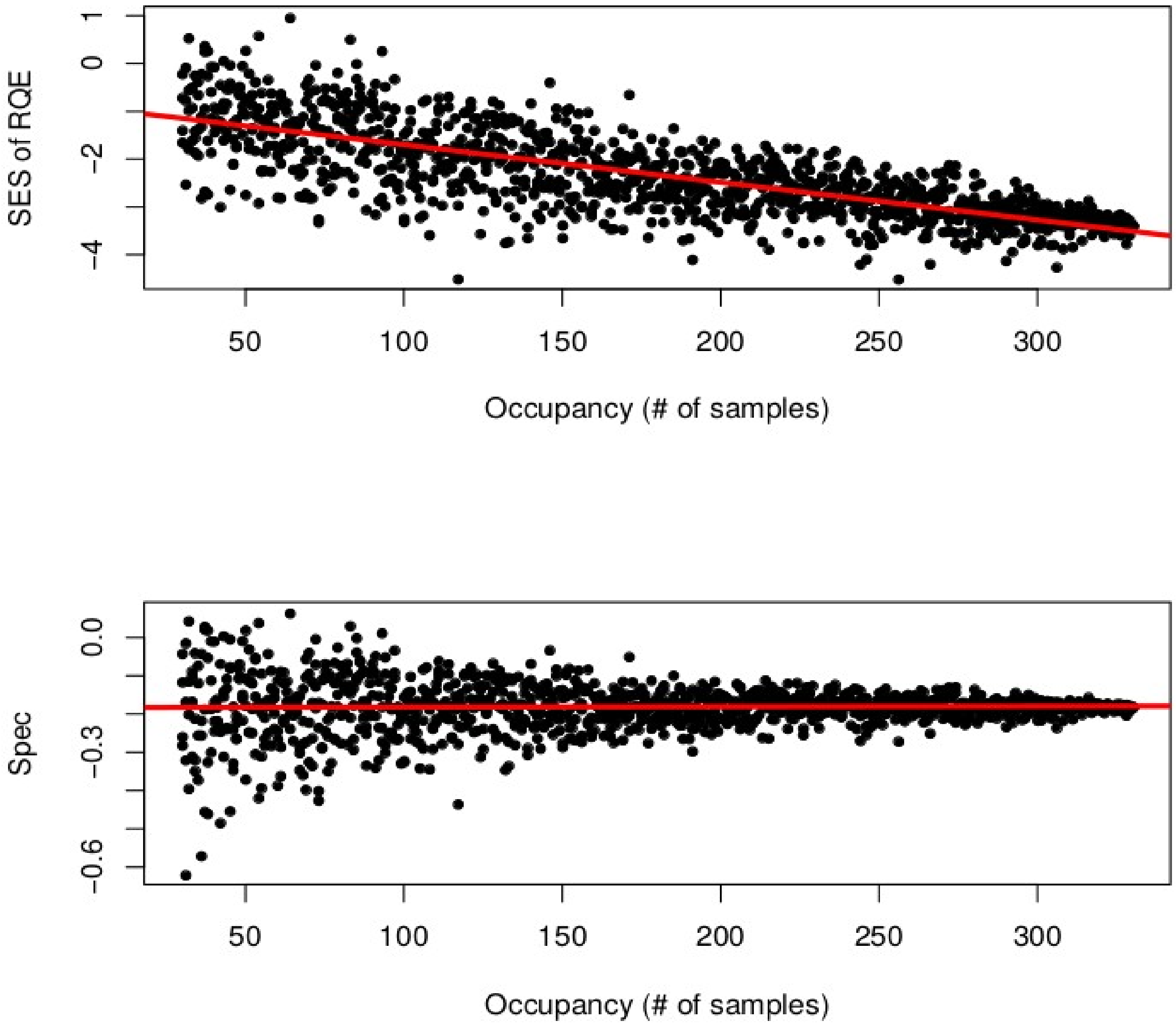


Supplementary Figure S5: Sensitivity of SES to occupancy (top) and sensitivity of *Spec* to occupancy (bntoom). Top: the standardized effect size (SES) of RQE (equation 1) was calculated using real microbial data (from the Hawaiian Endophytes data set) with elevation as the environmental variable. The number of samples where the species was present (occupancy) was lowered by randomly selecting samples and making the focal species absent in those samples. Counterintuitively, SES shows stronger specificity to species with higher occupancy, and weaker specificity when the species with lower occupancy. This is undesirable because a species with very strong specificity may be present in only a few samples. This occurs because SES is standardized using a distribution of values generated with permuted species weights. Higher occupancy means that the statistic (RQE, in this case) is more consistent across permutations, leading to a lower standard deviation. Lower occupancy reverses this trend, resulting in a more stochastic RQE calculation, leading to a higher standard deviation. Bottom: our statistic, *Spec*, is not sensitive to occupancy.

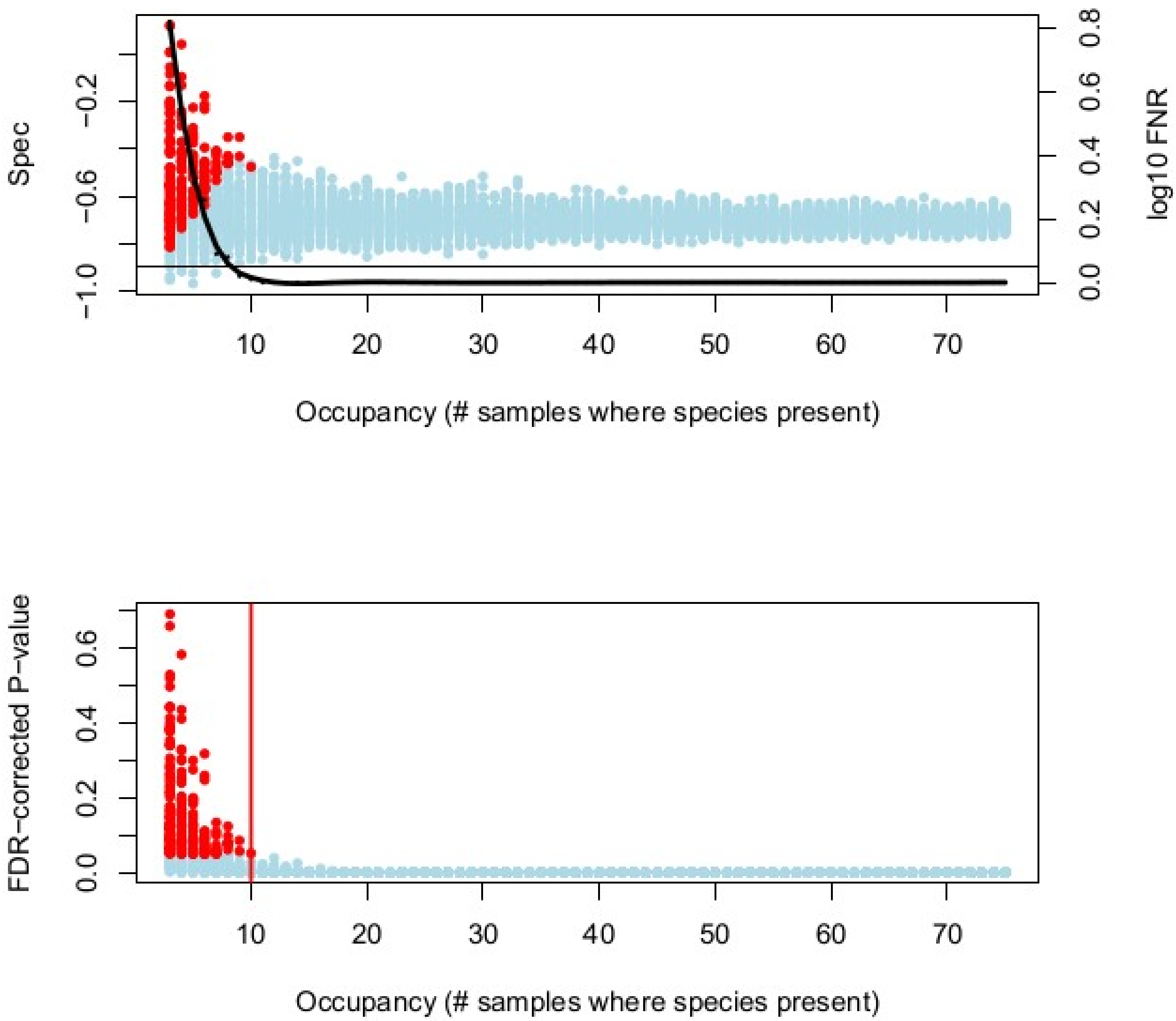


Supplementary Figure S6: Sensitivity of *Spec* to occupancy. For this analysis, a species observed in the Endophyte data set was subjected to random modification by selecting *n* data points within *p* (Equation 1) and replacing them with zeroes, for a range of values of *n*, including duplicates. Occupancy was calculated for each surrogate *p* vector produced this way, as the count of values within that vector above zero, and *Spec* was calculated for each, using elevation as an environmental variable. This analysis was conducted on more species than just the one shown in this figure (not shown), but the results were similar for each. False negative rate (FNR) was calculated for each unique value of *n* as the fraction of FDR-corrected *P*-values above 0.05, since the species vectors used in this analysis exhibited strong specificity before random modification as above. In the top panel, *Spec* is flat with respect to occupancy, although as occupancy is lowered, error in *Spec* increases, as expected. FNR was visualized by fitting a basis spline, shown as a black curve, and *P*=0.05 is shown as a horizontal black line. *P*-values are plotted alone in the bottom panel.

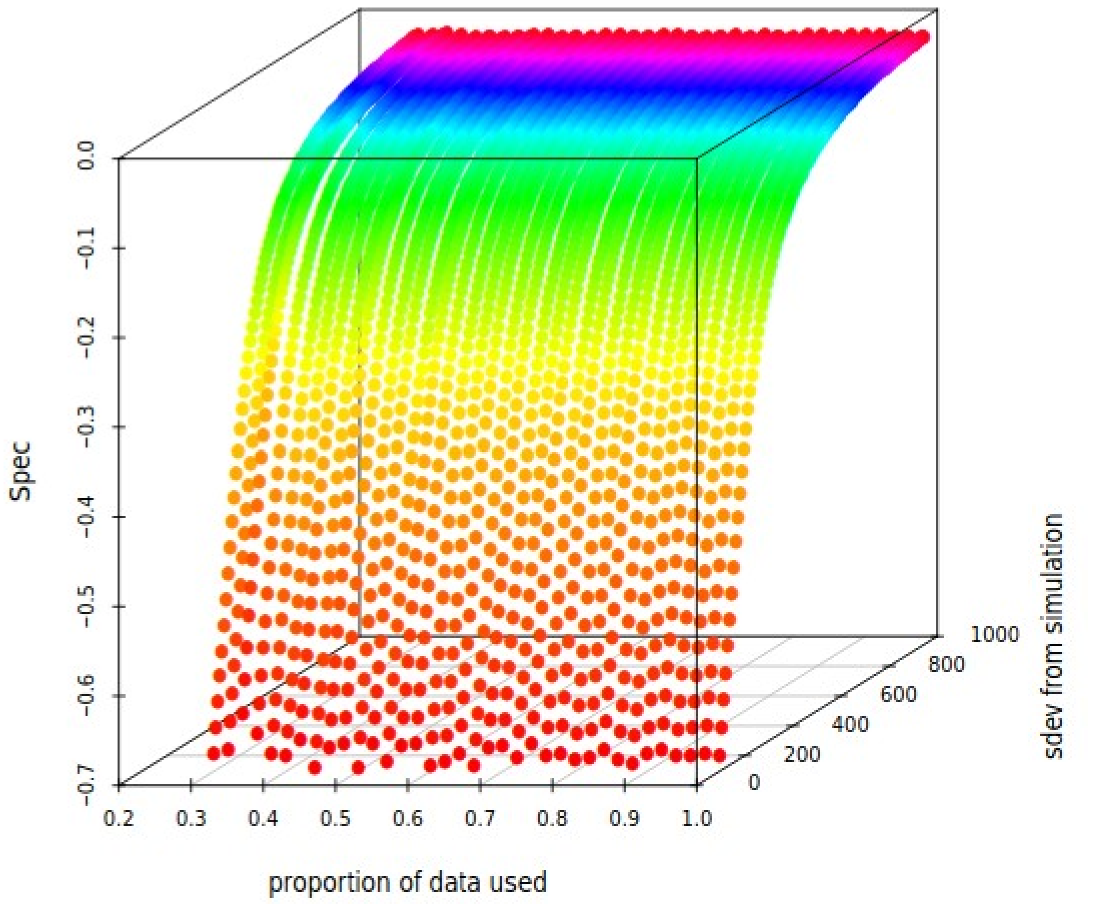


Supplementary Figure S7: Sensitivity of *Spec* to data set size. This figure shows a similar analysis to Supplementary Figure S3, except with an added dimension of data set size. The size of the data set was randomly downsampled by randomly eliminating samples (horizontal axis), from 500 samples (1.0) to 150 samples (0.3). Specificity was varied as before using SD (slanted axis), and *Spec* was calculated for each surrogate data set (vertical axis). The response of *Spec* to simulated specificity is flat with regard to the proportion of data used, showing that *Spec* is not sensitive to the size of a data set.

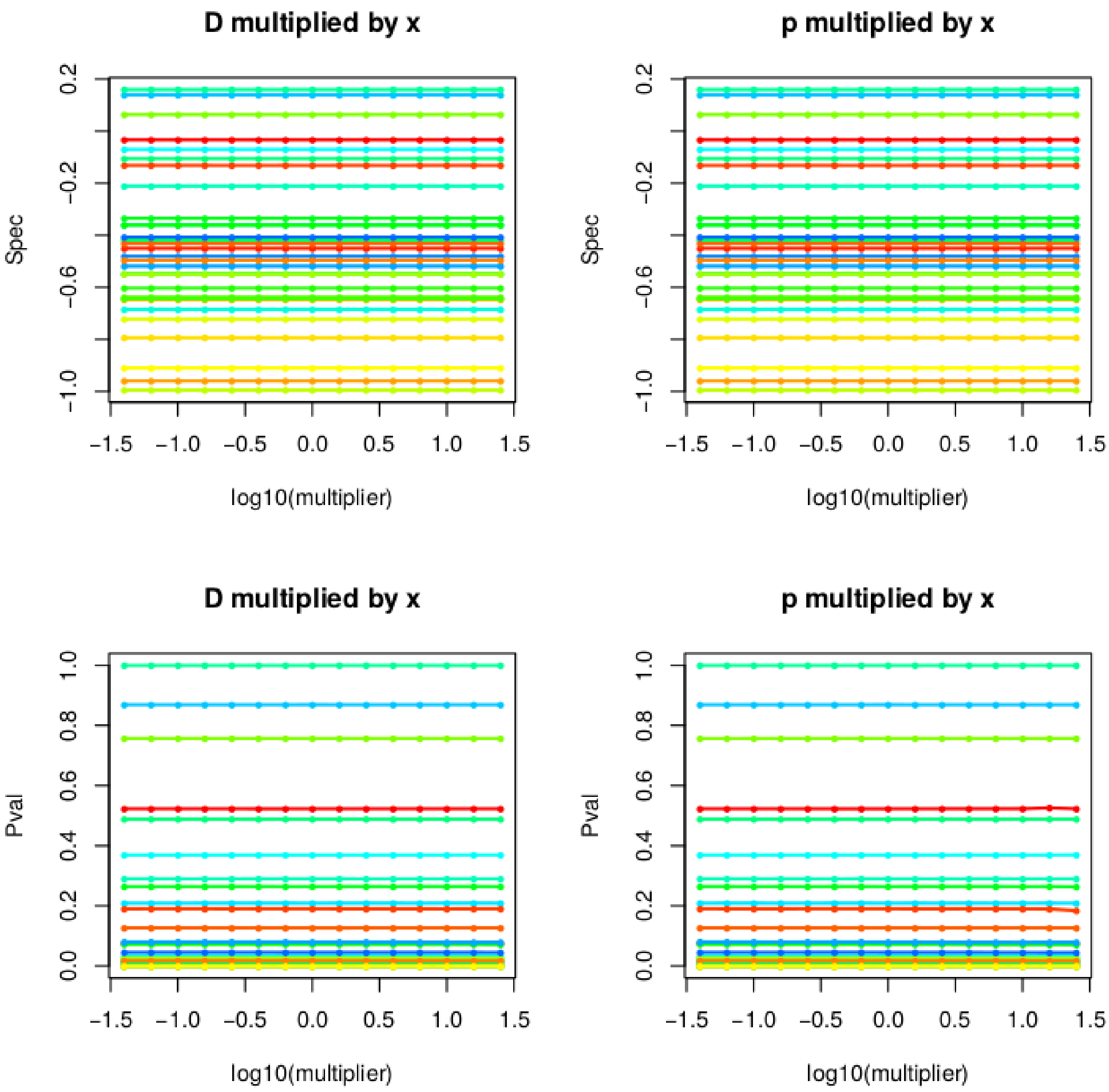


Supplementary Figure S8: Sensitivity of *Spec* to scaling of *p* and *D*. Both *p* and *D* can be scaled (multiplied by a scalar) and it will not change the output value of *Spec*, or the corresponding *P*-value. Separately, *p* and *D* inputs were multiplied by values ranging from $10^{-1.5}$ to $10^{1.5}$, and *Spec* was calculated using default settings, for 30 randomly selected OTUs from the Endophyte data set (each assigned a random color in this figure). In the subfigure titles, “x” refers to the x-axis variable, the range of values by which *p* or *D* was scaled. Note that before scaling, all $0 \leq p \leq 1$, and all $0 \leq D \leq 1$. Since Euclidean distances used to transform vector inputs to *D* inputs are not sensitive to addition of a constant, these results mean that temperature in degrees Celsius or degrees Fahrenheit could be used as environmental variables and the same results would be obtained.

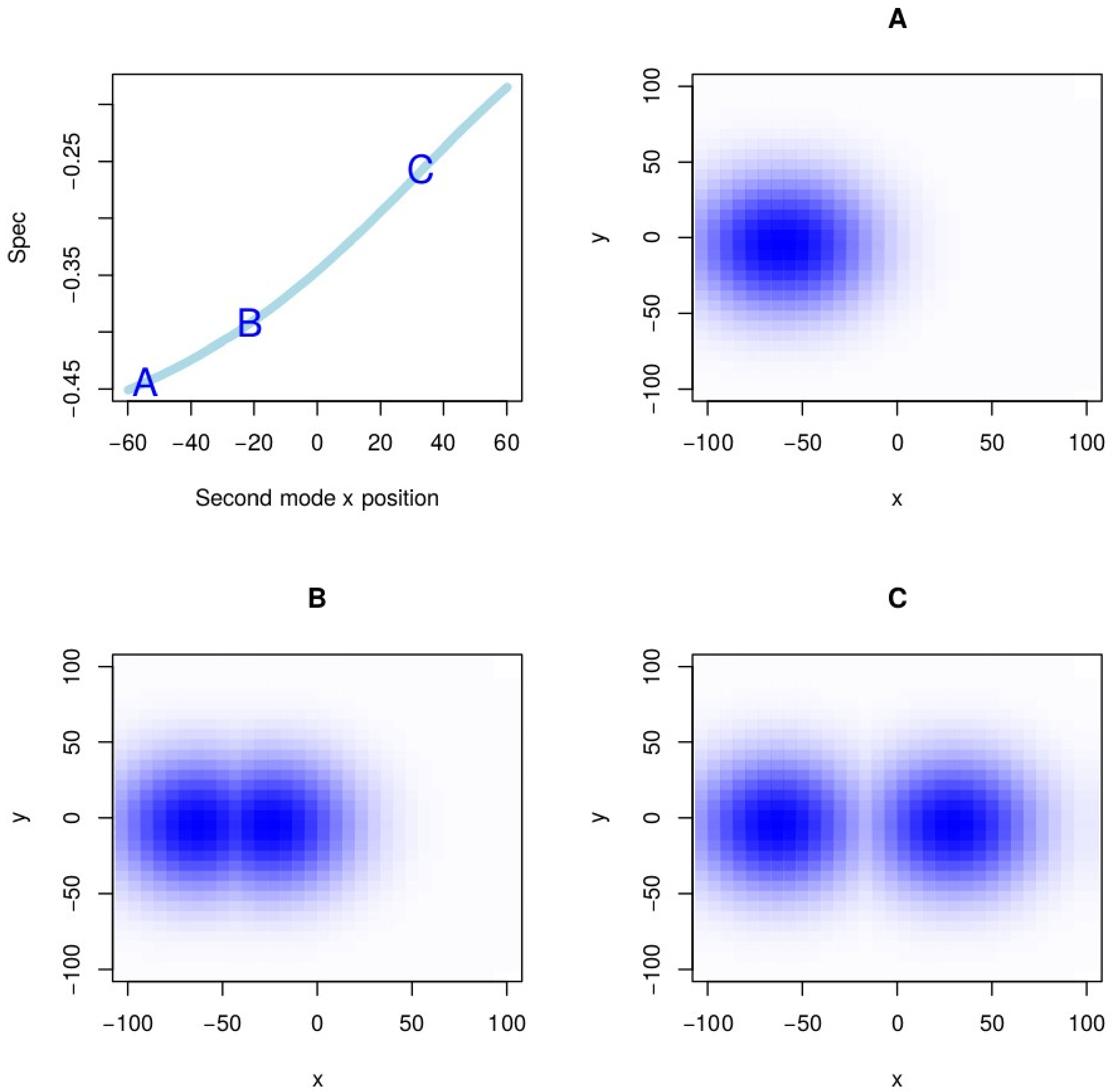


Supplementary Figure S9: Sensitivity of *Spec* to multimodality. A multimodal geographic distribution was simulated as an additive combination of bivariate normal distributions (x and y are geographic dimensions, e.g. longitude and latitude). When the two distributions had very close means (A), they simplify to a more unimodal distribution. The means were then moved farther apart to gradually make the distribution "more bimodal" (B, C). The results are plotted in the top-left plot, with labels corresponding to the distributions shown. This analysis shows that unimodal distributions give stronger *Spec* than bimodal distributions, and that this trend is continuous as modes are more distinct. However, multimodal specificity still fits our definition of generalized specificity, since the breadth of environment where the species is found is still reduced compared to random chance. This is shown by the consistent negative *Spec* values (and significant *P*-values, not shown).

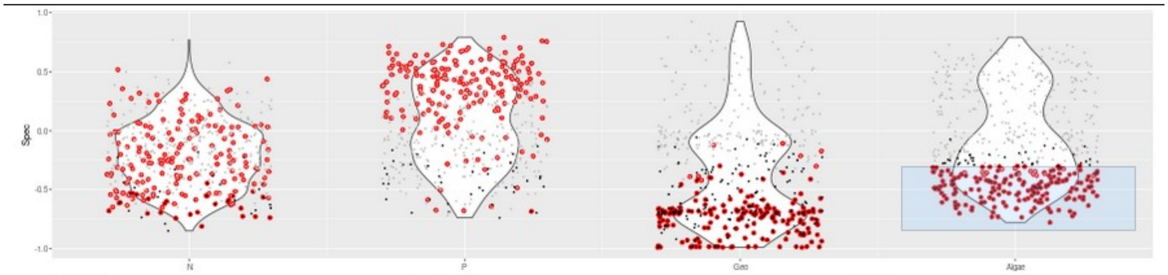


Search string:

Search field: FeatureID

Ignore Case? TRUE

P≤α color: #000000

P>α color: #BEBEBE

α: 0.05

Point size: 0.5

Point type: 19

Brush hilight color: #FF0000

Brush hilight size: 2

Download type: All data (long)

Download

Fields to show in brushed points (below):

☑ FeatureID ☐ N_Spec ☐ P_Spec ☐ Geo_Spec ☐ Algae_Spec ☐ N_Pval ☐ P_Pval ☐ Geo_Pval ☐ Algae_Pval ☐ eval ☐ hit ☐ k ☐ p ☑ c ☑ o ☑ f ☑ g ☑ s

Brushed points:

**Show** 10 **entries**

| | FeatureID | c | o | f | g | s |
|---|---|---|---|---|---|---|
| 3 | ASV0003 | Nostocophycideae | Nostocales | Nostocaceae | Nostoc | |
| 7 | ASV0007 | Synechococcophycideae | Pseudanabaenales | Pseudanabaenaceae | Leptolyngbya | |
| 11 | ASV0011 | Oscillatoriophycideae | Chroococcales | Xenococcaceae | | |
| 13 | ASV0014 | Chloroplast | Stramenopiles | | | |
| 31 | ASV0040 | Synechococcophycideae | Pseudanabaenales | Pseudanabaenaceae | | |
| 55 | ASV0071 | Cytophagia | Cytophagales | Cytophagaceae | Spirosoma | |
| 57 | ASV0073 | [Saprospirae] | [Saprospirales] | Chitinophagaceae | | |
| 65 | ASV0094 | [Saprospirae] | [Saprospirales] | Saprospiraceae | | |
| 67 | ASV0096 | Actinobacteria | Actinomycetales | Pseudonocardiaceae | Pseudonocardia | |
| 68 | ASV0097 | Sphingobacteriia | Sphingobacteriales | Sphingobacteriaceae | | |

Showing 1 to 10 of 192 entries — Previous 1 2 3 4 5 … 20 Next

Supplementary Figure S10: Screenshot of *specificity.shiny* visualization of Antarctic cryoconite hole data. This is an alternative visualization of the results presented in Figure 3, with jittered points for each species shown overlaying the violins. This enables species to be selected, as shown by the light blue box around species with strong specificity to algae. Those same species are present in other violins as well, and are shown outlined in red across the whole visualization. From this, we can see that species with strong specificity to algae also have strong geographic specificity, and weaker specificity to phosphorus (P) concentration. Taxonomy or other metadata for selected species is shown at bottom, and data shown can be filtered by these metadata as well (not shown).
